# A generalizable epigenetic clock captures aging in two nonhuman primates

**DOI:** 10.1101/2022.11.01.514719

**Authors:** Elisabeth A. Goldman, Kenneth L. Chiou, Marina M. Watowich, Arianne Mercer, Sierra N. Sams, Julie E. Horvath, Jordan A. Anderson, Cayo Biobank Research Unit, Jenny Tung, James P. Higham, Lauren J.N. Brent, Melween I. Martínez, Michael J. Montague, Michael L. Platt, Kirstin N. Sterner, Noah Snyder-Mackler

## Abstract

Epigenetic clocks generated from DNA methylation array data provide important insights into biological aging, disease susceptibility, and mortality risk. However, these clocks cannot be applied to high-throughput, sequence-based datasets more commonly used to study nonhuman animals. Here, we built a generalizable epigenetic clock using genome-wide DNA methylation data from 493 free-ranging rhesus macaques. Using a sliding-window approach that maximizes generalizability across datasets and species, this model predicted age with high accuracy (± 1.42 years) in held-out test samples, as well as in two independent test sets: rhesus macaques from a captive population (n=43) and wild baboons in Kenya (n=271). Our model can also be used to generate insight into the factors hypothesized to alter epigenetic aging, including social status and exposure to traumatic events. Our results thus provide a flexible tool for predicting age in other populations and species and illustrate how connecting behavioral data with the epigenetic clock can uncover social influences on biological age.

## INTRODUCTION

Chronological age is the predominant risk factor for most chronic, non-communicable diseases. However, chronological age *per se* cannot capture individual variation in health and disease risk beyond that which is associated with the passage of time. Measures of biological age aim to capture this variation to improve predictions of individual morbidity and mortality risk. The epigenetic clocks constructed independently by Horvath (2013) and Hannum and colleagues (2013) in humans were the first models of DNA methylation aging (also called epigenetic aging) to gain widespread use. Work in humans has found that accelerated epigenetic aging–when predicted biological age exceeds chronological age–is associated with increased risk of death (M. E. Levine et al., 2018; Marioni et al., 2015, 2016) and increased susceptibility to several hallmark diseases of aging (Ambatipudi et al., 2017; M. Levine et al., 2015; Wu et al., 2010; Yang et al., 2016; Zheng et al., 2016). Epigenetic clocks are also responsive to socio-environmental exposures associated with more rapid physiological and/or cognitive aging, such as trauma incurred during active military combat (Boks et al., 2015), childhood adversity (Austin et al., 2018; Marini et al., 2020; McCrory et al., 2022), and alcohol and tobacco use (Beach et al., 2015). However, the pathways through which social and environmental factors “get under the skin” to influence disparities in disease and mortality risk are difficult to investigate in humans alone. The development of epigenetic clocks in nonhuman animal models can help address this limitation by linking this promising tool to experimental studies, multigenerational field studies, and comparative analyses across species.

Rhesus macaques (*Macaca mulatta*) and baboons (*Papio* spp.) are excellent models for human aging because they are close evolutionary relatives for whom aging and survival are strongly dependent on characteristics of the physical and social environment (Blomquist et al., 2011; Chiou et al., 2020; Ellis et al., 2019). Such parallels are important because the impact of the environment on the progression of aging may not manifest similarly in short-lived models of mammalian aging, such as mice and rats. Indeed, questions that have been notoriously challenging to study in humans can often be addressed in these closely related nonhuman primates. For example, experimental life course studies in rhesus macaques have expanded our understanding of the causal effects of dietary restriction on life- and healthspan (Colman et al., 2014; Mattison et al., 2017). In addition, in social nonhuman primates, the association between social status and indicators of aging and life expectancy mirror some aspects of the social gradient of health in humans (Snyder-Mackler et al., 2020). Because nonhuman primate social systems are variable (Abbott et al., 2003), studying different social conditions and positions in the social hierarchy may point to specific variables that promote or detract from the impacts of extrinsic challenges on an individual’s health and survival.

The most extensively applied DNA methylation clocks to date (Hannum et al., 2013; Horvath, 2013) were built using DNA methylation data from humans generated on Illumina Infinium microarrays. By contrast, many recent studies of DNA methylation in nonhuman organisms have used high-throughput bisulfite sequencing (BS-seq) approaches (e.g., Chatterjee et al. 2013; Hahn et al. 2017; Chen et al. 2015; Pegoraro et al. 2016; Platt et al. 2015; Stubbs et al. 2017; Wang et al. 2017), for at least three reasons. First, while arrays designed for humans can sometimes be applied in other nonhuman primates (Hernando-Herraez et al., 2013; Ong et al., 2014), doing so requires stringent filtering for DNA sequence mismatches and changes in CpG site location, reducing the amount of usable data relative to studies in humans (Pichon et al., 2021; Teschendorff & Relton, 2018). Second, while designing species-specific arrays is also feasible, such tailored tools are often higher in cost. Third, many researchers have been interested not only in epigenetic aging, but also in patterns of differential methylation (e.g., age, genotype, or environment-related), which sometimes occur outside of the regions traditionally targeted by DNA methylation arrays (W. Zhang et al., 2015; Y. Zhang et al., 2016). Notably, this limitation also applies to recently developed multi-species arrays like the HorvathMammalMethylChip (Arneson et al., 2021). The multi-species array has provided extensive insight into epigenetic aging across many mammals (e.g., Horvath et al. 2022; Wilkinson et al. 2021) while capturing DNA methylation at only 38,000 highly conserved, non-randomly distributed CpG sites. Consequently, generalizable tools to study epigenetic aging using BS-seq data are also needed. Here, we developed the RheMacAge model, a nonhuman primate epigenetic clock that facilitates comparisons among Old World monkey species often used as models for human aging. We first developed a generalizable epigenetic clock model that can predict chronological age using BS-seq data with high accuracy. Next, we applied our model to two independently generated datasets to demonstrate not only the *cross-study* but *cross-species* applicability of our approach in two nonhuman primates with exceptional research importance, rhesus macaques and baboons. Finally, we used the model to test whether social status or exposure to an adverse climate event were associated with variation in epigenetic aging.

## MATERIALS AND METHODS

### Study Population and Sample Collection

Our primary dataset consisted of 563 whole blood samples (number of unique individuals=493) collected from rhesus macaques living on Cayo Santiago, an island 1 km off the coast of Puerto Rico that is home to over 1,800 free-ranging rhesus macaques. The population is managed by the Caribbean Primate Research Center (CPRC) as part of a long-term field station in operation since 1938. The macaques are provisioned with food and water but are otherwise allowed to roam freely, self-organize into social groups, and do not face the threat of predation (Rawlins & Kessler, 1986). Collectively, these conditions provide opportunities to investigate how interactions between ecology and behavior influence environmentally-responsive molecular mechanisms (such as DNA methylation) in the absence of other confounding factors (e.g., variable nutrition and predation).

The data in this study were collected from 273 female and 220 male rhesus macaques, aged 1.44 months to 28.82 years (birth dates recorded by CPRC census takers). We collected blood samples and behavioral data on these animals between 2010 to 2018 (with the exception of 2017, when sampling was not possible due to Hurricane Maria). On Cayo Santiago, the average age at sexual maturity for a female is 4 years, and median lifespan for an adult female is 18 years (Chiou et al., 2020). By comparison, human females in the United States reach sexual maturity at an average age of 14 years (Susman et al., 2010) and have a median lifespan of 80.5 years (National Center for Health Statistics 2021).

Sixty-six individuals were sampled more than once during this study (see **Supplemental File 2** for detailed metadata). Blood was collected into K3 EDTA vacutainer tubes (BD Biosciences, Franklin Lakes, NJ) and placed on ice until storage at −80°C (within 8 hours of collection). All work was reviewed and approved by the Institutional Animal Care and Use Committees of the University of Washington (assurance number A3464-01) and the University of Puerto Rico, Medical Sciences Campus (assurance number A4001-17). This work also adheres to the American Society of Primatologists Principles for the Ethical Treatment of Nonhuman Primates.

### RRBS Data Generation

Genomic DNA was isolated from whole blood using the Qiagen Blood and Tissue DNA kit (QIAGEN, Hilden, Germany). To measure CpG methylation, we used Reduced Representation Bisulfite Sequencing (RRBS), a version of bisulfite sequencing (BS-seq) that uses an initial restriction enzyme digest to concentrate sequencing in and near CpG-rich regions of the genome (Gu et al., 2011; Meissner, 2005).To prepare RRBS libraries, we followed the library preparation protocol detailed on the Snyder-Mackler Lab website (https://smack-lab.com/wp-content/uploads/2020/03/SMack_Lab_RRBS-with-Zymo-EZDNA-MagBead.pdf). Briefly, we digested extracted DNA using the *Msp1* restriction enzyme, which cuts at CCGG sites, ligated NEBNext methylated adapters (Illumina Inc., San Diego, CA), bisulfite converted the DNA using the Zymo EZDNA Methylation-Lightning™ Kit (Zymo Research, Irvine, CA), and PCR-amplified the final fragments with unique dual molecular indices for each sample. RRBS libraries were sequenced in two batches. Batch 1 contained 104 samples (2×50bp reads) sequenced on an Illumina NovaSeq S2 flowcell. Batch 2 was made up of 527 samples (2×100bp reads) and sequenced on a NovaSeq S4 flowcell.

### Alignment and Preprocessing

We trimmed adapter and low-quality sequences with TrimGalore! (v0.4.5) (Martin, 2011). We then aligned trimmed reads to the *in silico* bisulfite-converted reference genome (Mmul10) using the default settings in Bismark (v0.20.0) (Krueger & Andrews, 2011) for all but two parameters (--score-min and -R; see the **Methods Supplement**). We parallelized sample aggregation with GNU parallel (Tange, 2018) and BedTools (v2.24.0) (Quinlan & Hall, 2010). Unless otherwise stated, all subsequent analyses were carried out in RStudio (v1.4.1106) (RStudio Team, 2015).

### Development of a Generalizable Epigenetic Clock

#### Site-based modeling approach

To limit the inclusion of invariant or uninformative sites, we first removed CpG sites with missing data in more than 10% of the training samples and samples missing more than 25% of CpG sites in the filtered dataset. Next, we removed constitutively hypo- or hypermethylated sites (those with median percent methylation less than 10% or greater than 90% across samples), and sites with less than 5X median coverage, resulting in 196,345 CpG sites in the site-based dataset. A detailed description of sample filtering criteria can be found in the **Methods Supplement**.

Next, we imputed missing and low coverage (< 5X) sites in the 196,345 CpG site dataset using BoostMe (v0.1.0) (L. S. Zou et al., 2018). On average, 12.5% of sites were imputed per sample. After removing sites mapping to sex chromosomes and those containing one or more inadmissible values (non-real numbers that result from when BoostMe attempts to divide by zero), the dataset contained 185,153 unique CpG sites.

#### Sliding window-based modeling approach

To improve model generalizability, we compared the performance of a traditional site-based model to a 1 Kb, non-overlapping sliding-window based approach that we reasoned might capture more shared loci across samples and datasets. In each 1 Kb window of the genome, we calculated the percent methylation for a given sample as the number of reads covering methylated CpG sites, divided by the total number of reads covering CpG sites in that window. We implemented the same filtering strategy described above by excluding windows that had missing data in more than 10% of samples (leaving 279,052 windows), samples that were missing data for more than 25% of windows in the dataset, windows that were constitutively hypo- or hypermethylated, and those with less than 5X median coverage, leaving a final set of 161,289 windows.

Missing values and those with < 5X coverage at a given site were imputed using BoostMe. The average proportion of imputed windows was 1.98% per sample and ranged from 0.05% to 23.2%. Notably, the average proportion of features imputed for the window-based approach was six-fold lower than that for the site-based dataset (1.98% as compared to 12.5%). Following imputation, removal of windows containing non-real numbers, and removal of those mapping to sex chromosomes, 159,472 windows were retained for calibrating the epigenetic clock.

All three datasets used to train, validate, and/or test the model were processed in an identical manner.

### Model Training and Optimization

We then generated age-prediction models independently for the site-based and window-based datasets. We used elastic net regression implemented using the R package glmnet (v4.1-1) (Friedman et al., 2010) and leave-one-out cross validation (LOOCV) to train our model. To perform LOOCV, one sample was held out at a time, and ‘proto-models’ were generated on the remaining N-1 samples using 10-fold internal cross validation. The “best” proto-model (the proto-model with the lowest mean absolute error) was then used to predict the age of the held-out sample. This process was repeated for each sample in the dataset. Once all 563 age predictions were generated, we regressed all predicted age values onto known chronological ages to evaluate the predictive performance of the dataset in our primary dataset.

We defined a measure of age acceleration, termed “residual epigenetic age” (similar to Horvath’s [2013] “delta age”) by taking the residuals from a loess (locally estimated scatterplot smoothing; span = 0.75) regression of predicted onto chronological age to identify individuals who appear to be aging more (or less) rapidly than expected given their chronological age. By taking the residuals from a loess regression, we can detect potentially meaningful deviations from the expected rate of aging while accounting for systemic effects of the model (e.g., the influence of chronological age) and non-linear pace of epigenetic aging. We adjusted for a marginally significant effect of sex on epigenetic age (*p* = 0.05) by calculating residual epigenetic age for females and males separately.

### Model Validation in Independent Datasets

We sought to test the generalizability and performance of both the site- and window-based models (“Cayo” models) using two independently generated RRBS datasets (“Yerkes” and “baboon” datasets). However, the site-based model’s applicability was constrained by the limited overlap between the site-based RRBS independent datasets (**Figure S1**).

To test the generalizability of the window-based model to populations and study systems of the same species, we used samples from 43 female rhesus macaques, aged 3.1 to 20.1 years, housed at Yerkes National Primate Research Center. RRBS libraries were generated from purified classical monocytes (CD3^-^/CD14^+^) collected in an unrelated study examining dominance rank effects on gene regulation and immune function (Snyder-Mackler et al., 2016). Second, to test generalizability in another important model for human aging, we applied our model to a second RRBS dataset generated from 271 whole blood samples collected from wild baboons living in Amboseli National Park in Kenya (Anderson et al., 2021) (SRA project accession PRJNA648767). This dataset contains samples collected from 138 females and 133 males, aged 1.93 to 26.34 years. The data processing workflow was identical across the Cayo, Yerkes, and baboon datasets.

To generate a single epigenetic clock for predicting age in independent datasets, we quantile normalized methylation values across features and across samples independently for the Yerkes macaque and Amboseli baboon datasets. Next, we determined optimal hyperparameter settings using the caret package (v6.0-86) (Kuhn, 2019, https://topepo.github.io/caret/) by performing a grid search across two hundred combinations of alpha (the parameter that controls how similar the model is to lasso versus ridge regression ((H. Zou & Hastie, 2005) and lambda (the regularization parameter) using repeated (3x) 10-fold cross validation on all 563 Cayo samples. After LOOCV models were generated, we performed an elastic net regression with the optimized hyperparameter values on the entire dataset and applied the resulting model to the two external datasets (Yerkes and baboon).

### Effects of Social and Environmental Adversity on Epigenetic Aging

To explore how heterogeneity in the social environment is reflected in biological aging, we tested the relationship between dominance rank and epigenetic aging in the Cayo Santiago rhesus macaque samples. First, we quantified dominance rank for 81 males and 116 females from dyadic win-loss interactions between individuals within a social group using behavioral data collected prior to each blood draw. The dominance rank of an individual represents the proportion of same-sex individuals in their group that they out-ranked. In 2010-2014, the trap and release season was Jan-Mar, and rank data were calculated from behavioral data recorded in April-Dec of the previous year. In late 2014 until 2018, the trap and release season was moved to Oct–Dec. For samples collected in this time period, rank data were thus calculated from behavioral data recorded in Jan–Oct of the same calendar year. Next, we tested if dominance rank is associated with epigenetic age by modeling residual epigenetic age as a function of dominance rank for females (aged 6.01 – 27.93 years) and males (aged 5.89 – 22.78 years) separately. Additionally, we tested whether residual epigenetic age was driven by length of tenure in the social group among males in a larger subset of the Cayo Santiago population (n = 230, aged 3.86– 21.74 years). Rhesus macaque males attain their dominance largely via queueing instead of direct contest such that longer residency times predict higher rank (Kimock et al., 2019, 2022). Tenure length is thus a useful proxy of male rank and is advantageous as a measure because demographic records have been collected over a longer period of time compared to behavioral data.

Finally, motivated by the observation that Cayo Santiago macaques that survived Hurricane Maria in 2017 exhibited “aged” blood transcriptomes (Watowich et al., 2022), we tested if Hurricane Maria had left a similar mark in epigenetic age. To do so, we modeled residual epigenetic age as a function of exposure to Hurricane Maria for all adult animals (aged at least 4 years). We excluded infants and juveniles due to previously observed differences in the rate of aging between human adults versus children and adolescents (Horvath, 2013).

## RESULTS

### RheMacAge *is a generalizable epigenetic clock that accurately predicts chronological age*

Although our site-based model was able to accurately predict age (Pearson’s r = 0.82 MAD = 2.11 years) **(Figure 1A),** it was significantly outperformed by the window-based model (Pearson’s r = 0.9, MAD = 1.42 years; 0.69-year difference in MAD between two models, *p_t-test_* = 5.36 x 10^-8^) (**Figure 1B**). The window-based model performed equally well in males and females (*p_t-test_* = 0.71). While both models predicted chronological age well, their predictions did not scale linearly with chronological age. Age predictions plateau at older ages (> 20 years among the macaques in our Cayo sample), as has been reported in other species, including humans (e.g., Levine et al. 2020; Horvath 2013).

**Figure 1.**
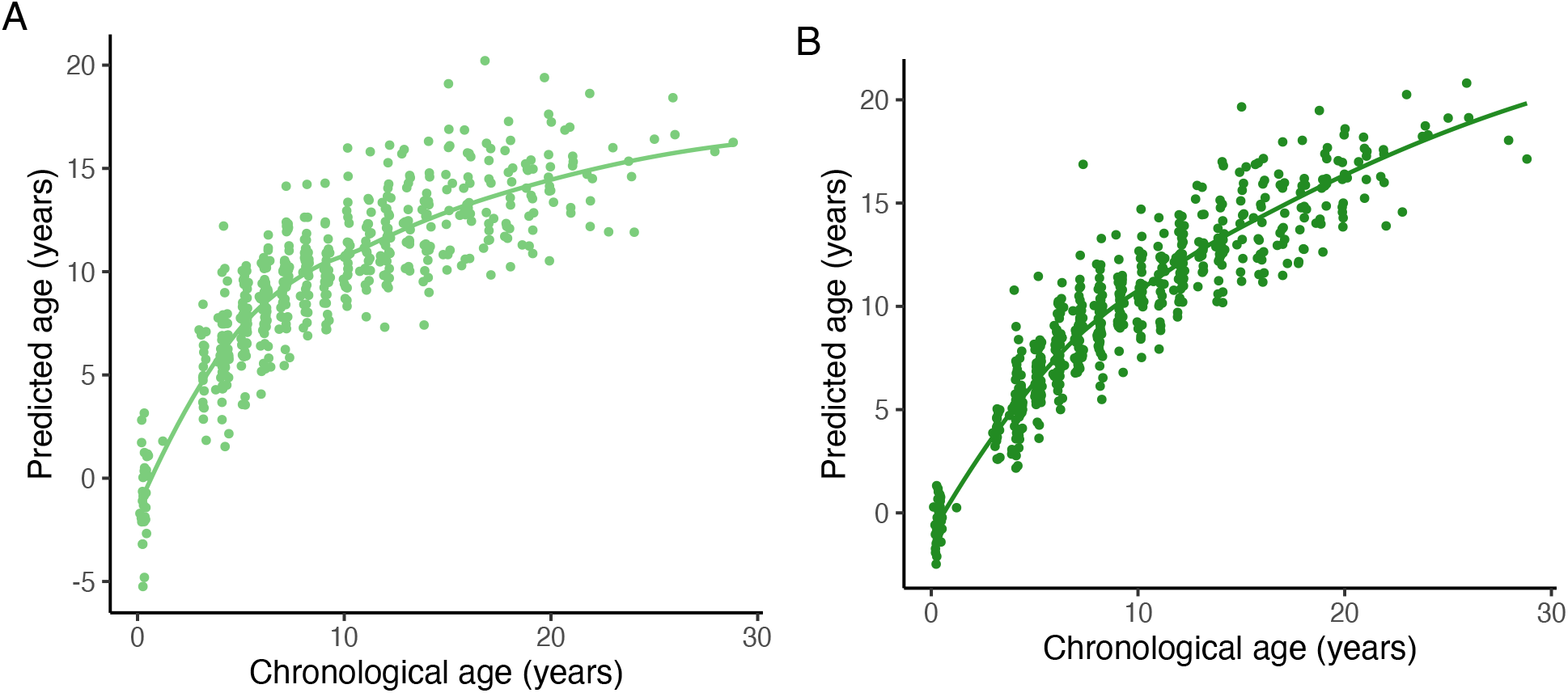
Comparison of Site-Based and Sliding Window-Based Models. **(A) Site-based model of methylation age successfully predicts known chronological age.** Known chronological age is highly correlated with epigenetic age predictions from our site-based epigenetic clock (Pearson’s r = 0.82, median absolute deviation between predicted and chronological age [MAD] = 2.11 years). Methylation data used to generate the site- and window-based clocks are from whole blood samples from a cross-sectional sample of rhesus macaques living on the island of Cayo Santiago (n samples = 549; n unique females = 267, n unique males = 217). Curved line shows line of best fit from univariate loess regression. **(B) Window-based model of methylation age successfully predicts known chronological age and outperforms the site-based model.** Known chronological age is more highly correlated with epigenetic age predictions in the window-based epigenetic clock (Pearson’s r = 0.9, MAD = 1.42 years) than the site-based clock. The model was generated using whole blood samples from rhesus macaque living on the island of Cayo Santiago (n samples = 563; n unique females = 273, n unique males = 220). Curved line shows line of best fit from univariate loess regression.

Although the site- and window-based datasets contained a similar number of loci after filtering within the Cayo data (~180K and ~160K, respectively), when applying these models to the external Yerkes data set, the overlap in features retained in both datasets was much higher when we used the window-based approach. Of the 185,153 CpG sites in the filtered Cayo dataset, 38% (70,439) of sites also passed the same filters in the Yerkes dataset, compared to 97% (155,347) of shared features for the window-based dataset (**Figure S1).** Given that the window-based model also showed superior performance in predicting age in the Cayo population, we used the window-based RheMacAge clock for all subsequent analyses.

The RheMacAge clock included 359 windows, of which 164 decreased in methylation with age (“hypomethylated windows”) and 195 increased in methylation with age (“hypermethylated windows”). CpG sites that exhibit age-dependent changes in methylation are often found in evolutionarily conserved regions of the genome (Mozhui & Pandey, 2017), and certain age-dependent patterns of methylation change are conserved between humans and mice (Spiers et al., 2016; Stubbs et al., 2017). Indeed, windows in the RheMacAge clock were modestly but significantly more evolutionarily conserved than windows that were not part of the clock (D = 0.09, *p* = 0.007, two-sample Kolmogorov-Smirnov test) (**Figures S2, S3**).

Since we collected repeated samples from 66 individuals, we were able to test if our model accurately tracked aging within an individual. For these 66 individuals (n=70 paired samples, because 4 individuals were sampled three times), 88.6% (62/70) exhibited increased biological age with increased chronological age (**Figure 2**). Thus, samples collected later in time were consistently predicted to be older than those collected at earlier points in time *(p* = 9.13 x 10^-12^, one-sided exact binomial test).

**Figure 2.**
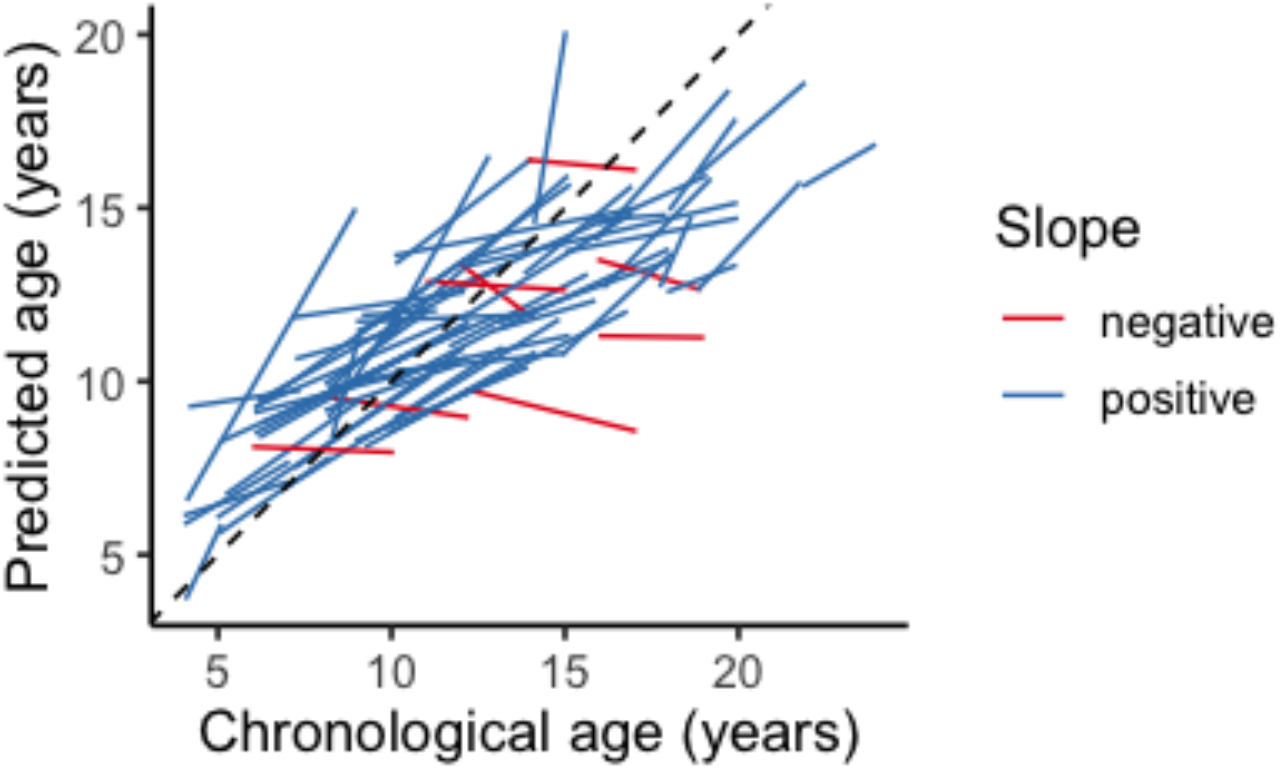
Predicted Aging Trajectories for Repeatedly Sampled Individuals. Predicted age increased over time in repeatedly sampled individuals (n = 66 individuals). 88.6% of epigenetic age predictions were higher in the sample collected when the individual was older (blue lines; 62/70 predictions; p = 9.13 x 10^-12^, one-sided exact binomial test). Dashed line shows x=y.

### The RheMacAge clock predicts age in two independent datasets, demonstrating cross-study and cross-species applicability

We evaluated our model (“Cayo”) in an independent macaque sample comprised of 43 female rhesus macaques from the “Yerkes” dataset (aged 3.1 to 20.1 years). DNA methylation-based age predictions were significantly correlated with chronological age (**Figure 3**) (Pearson’s r = 0.69, *p* = 2.65 x 10^-7^), with an MAD of 2.09 years. This suggests the RheMacAge clock can be generalized to predict age in rhesus macaque data generated from different populations and different cell types (monocytes versus all white blood cells), and by different laboratories.

**Figure 3.**
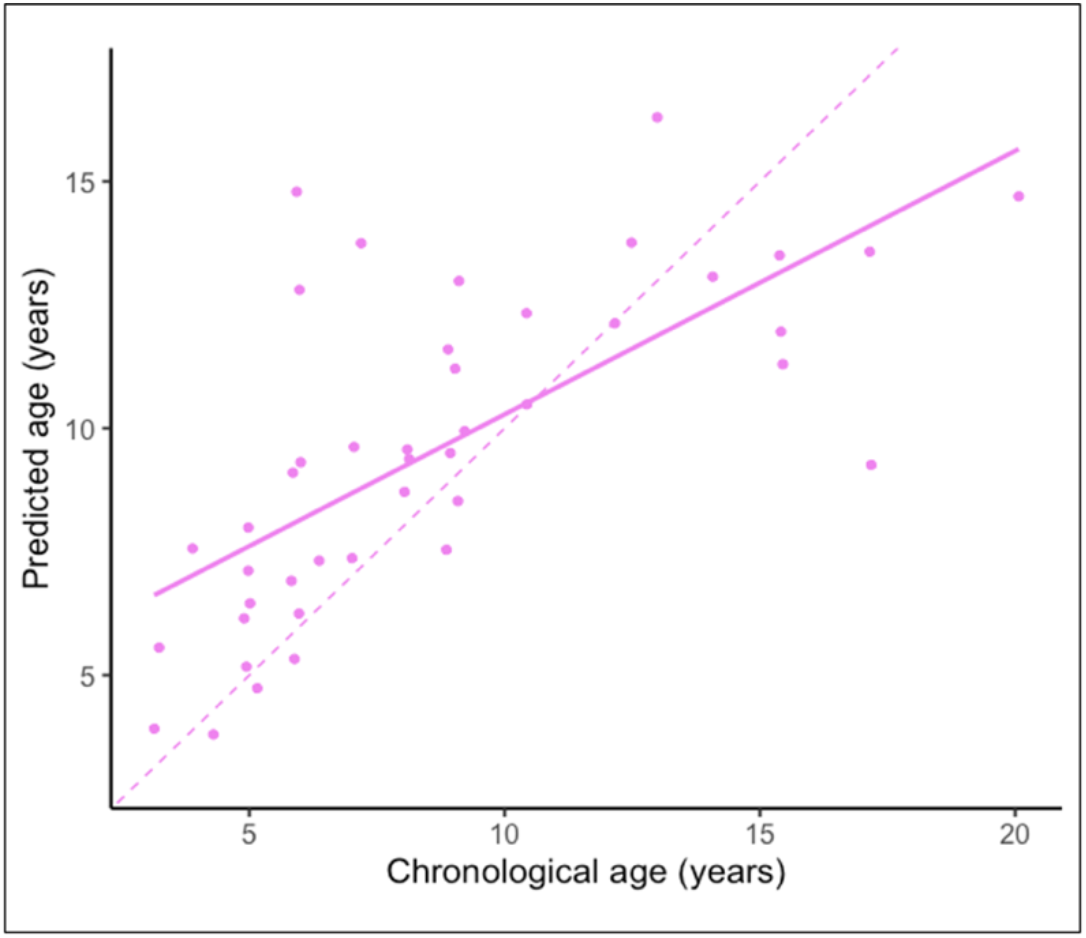
Predicted DNA Methylation Age for Yerkes Macaques Using RheMacAge Model. Predicted DNA methylation age for Yerkes rhesus macaques is correlated with chronological age (r = 0.69, MAD = 2.09 years). Solid line shows line of best fit from a univariate linear regression of predicted age onto chronological age.

When applied to individuals from our “baboon” dataset (aged 1.93 to 26.34 years), the RheMacAge clock was also able to predict chronological age surprisingly well, producing the same sex-specific patterns in the rate of aging described in previous work (Anderson et al., 2021). Specifically, prediction was better for male baboons than female baboons, especially at older ages (male Pearson’s r = 0.8, *p* < 2.2 x 10^-16^; MAD = 1.34 years, **Figure 4A**; female Pearson’s r = 0.74, *p* < 2.2 x 10^-16^; MAD = 2.19 years, **Figure 4B**). The RheMacAge clock predictions for baboons were not as accurate as a baboon-specific model (baboon clock: male MAD = 0.85 years, female MAD = 1.6 years). Importantly, the RheMacAge clock captured a similar biological signal to the baboon clock: residual epigenetic ages calculated from both models were significantly positively correlated for both sexes (males: r = 0.55, *p* = 5.28 x 10^-12^; females: r = 0.41, *p* =7.49 x 10^-7^; **Figures 4C** and **4D**). In addition, for male baboons, estimates of residual epigenetic age generated using the RheMacAge clock replicated the significant association (first reported in Anderson et al., 2021) between high social status and older epigenetic age (Pearson’s r = −0.47, *p* = 4.05 x 10^-7^, n = 104; the correlation is negative because lower values on an ordinal rank measure represent higher status). We found no significant association between rank and epigenetic age in female baboons.

**Figure 4.**
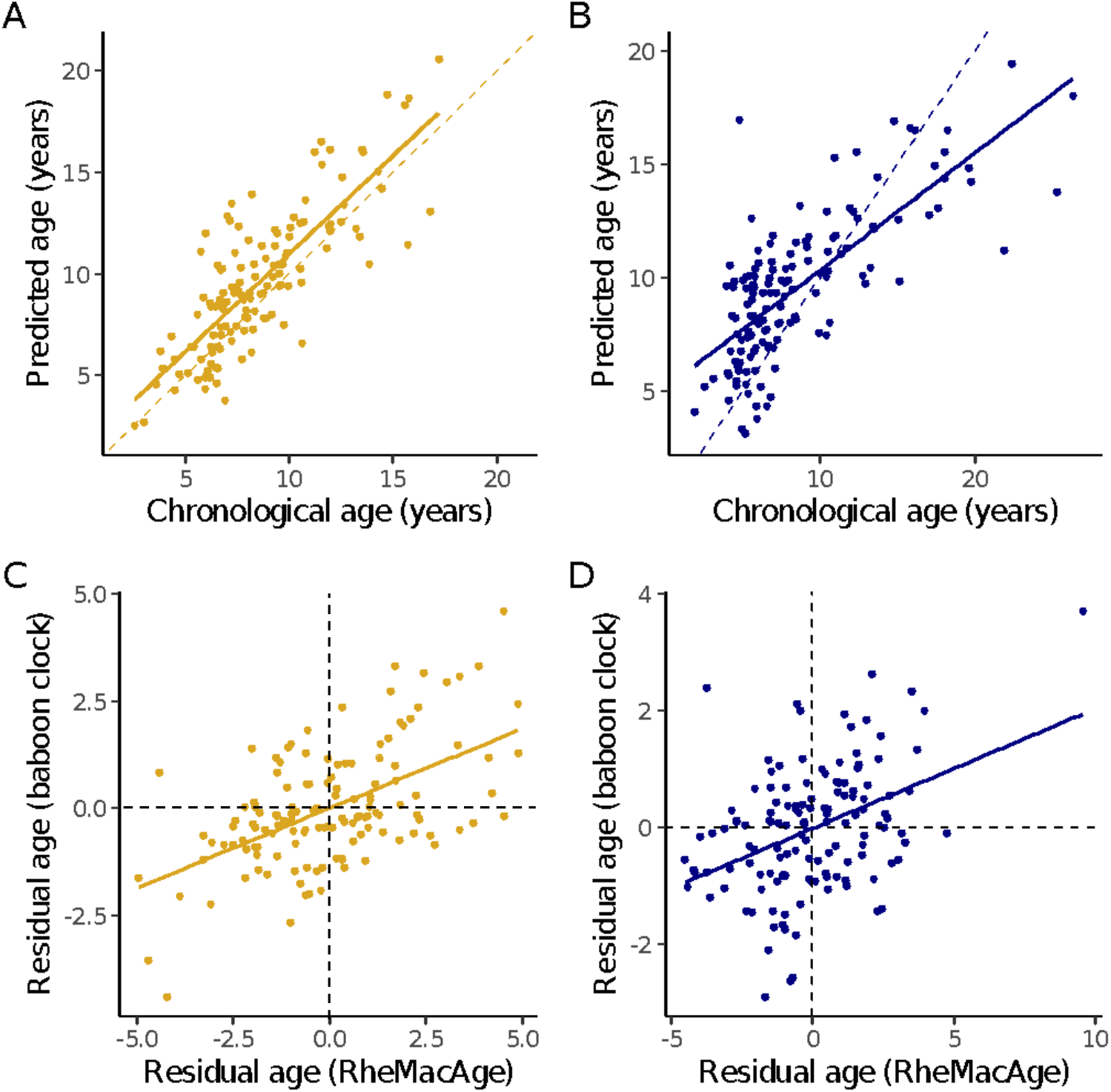
Predicted DNA Methylation Age for Amboseli Baboons Using RheMacAge Model. RheMacAge successfully predicts interspecies epigenetic ages from baboon DNA methylation data and recapitulates results from a baboon-specific methylation clock. Predicted age for **(A)** male and **(B)** female baboons using the RheMacAge epigenetic clock are highly correlated with known chronological age (males: r = 0.8, MAD = 1.34 years; females: r = 0.74, MAD = 2.19 years). The solid line shows the line of best fit from univariate linear regression of predicted onto chronological age. Dashed line shows x = y. Residual epigenetic age from the RheMacAge (x-axis) recapitulates residual ages from a baboon-specific clock (y-axis) for **(C)** males (r = 0.55, p = 5.28 x 10^-12^) and **(D)** females (r = 0.41, p = 7.49 x 10^-7^). Points in the bottom left (decelerated ages) and top right (accelerated ages) quadrants reflect concordance in residual epigenetic ages between the two clocks. Discordant predictions between the two clocks (i.e., one clock predicts accelerated age while the other predicts decelerated rate of aging) are in the top left and bottom right quadrants.

### Epigenetic age predictions for the Cayo macaques showed no association with social and environmental variables tested

We identified no association between residual epigenetic age and dominance rank in adult male (n = 60, β = −0.22, *p* = 0.3) or female (n = 83, β = −0.05, *p* = 0.77) rhesus macaques from Cayo Santiago. Residual epigenetic age in males also did not correlate with group tenure length (n = 59, β = −0.23, *p* = 0.33). Finally, contrary to our prediction, we found no relationship between hurricane exposure and residual epigenetic age (*p* = 0.24).

## DISCUSSION

### The RheMacAge clock accurately captures chronological and biological aging in two nonhuman primate models for human aging and offers a generalizable approach that can be used when developing epigenetic clock models from BS-seq data

Rhesus macaques and baboons are important biomedical models for human aging (Chiou et al., 2020; Huber et al., 2020). However, there are comparatively fewer ‘omics’ resources available for either species than for humans or mice (see Meer et al., 2018). To partially address this gap, here we present genome-wide DNA methylation data from 563 rhesus macaque samples along with new analyses of previously published data from 43 rhesus macaques and 271 baboons. The present study thus represents the largest study of DNA methylation carried out to date in nonhuman primates.

Our RheMacAge clock produced accurate age estimates from blood in an independent sample of captive female rhesus macaques and a large sample of wild baboons. We also observed a more rapid rate of epigenetic change with age in males versus female baboons that is consistent with results from Anderson et al. (2021), demonstrating that our approach can capture similar aging signatures across species. As such, the model serves as a useful biomarker of aging in blood that can facilitate research on the causes and consequences of the aging process. While the RheMacAge clock was only tested in two species, it may be suitable for application other primates (e.g., long-tailed macaques, sooty mangabeys) that are closely related to rhesus macaques and baboons. In addition, we offer a more generalizable approach that can be used when developing epigenetic clock models from BS-seq data more commonly generated from nonhuman animals.

### Species-specific socioecology is reflected in variation in epigenetic aging in baboons and macaques

We found no significant relationship between epigenetic aging and dominance rank in female or male rhesus macaques. This result differs from the effect described in male baboons, where high-ranking males exhibited older relative epigenetic ages compared to lower-ranking males (Anderson et al., 2021). On the surface, these findings appear discordant, but they may be a legitimate reflection of differences in dominance rank acquisition between rhesus macaques and baboons. Specifically, in the Amboseli baboons, male rank is determined by competitive interactions and mating occurs throughout the year. Maintaining alpha status requires significant energy expenditure during mate guarding and competition with other males, and alpha males have the highest glucocorticoid levels across the male status hierarchy (Alberts et al., 2006; Gesquiere et al., 2011). In contrast, male rhesus macaques obtain high social status through a less physically competitive queueing system in which males rise in rank as their tenure in a social group lengthens (Berard, 1999). Rhesus macaques are also seasonal breeders, meaning that direct male investment in mating effort is restricted to several months of the year (Bercovitch, 1997). Consequently, high status is not as difficult or energetically expensive to maintain in male rhesus macaques as it is in male baboons, and status in rhesus males is more predictable. Together, these species differences in the male competitive regime predict that high rank in male baboons is more energetically taxing, while high rank in rhesus males may be less stressful and/or demanding compared to lower ranking positions (Sapolsky, 2005). These differences point to strategies of mitigating distinct types of social stress as one mechanism that may be involved in dictating the rate at which the biological clock ‘ticks’.

Surprisingly, although exposure to Hurricane Maria was associated with accelerated immunological aging in the Cayo rhesus macaque transcriptome (Watowich et al., 2022), it was not significantly associated with residual epigenetic age in our study. One possible explanation for these observations is that the mechanisms underlying the clock are distinct from those that regulate age-related changes in the immune response (see Bell et al., 2019 for discussion of mechanisms). Further, the extent and velocity at which DNA methylation changes at sites typically captured in epigenetic clocks following exposure to adverse events is not known. Future studies will help enhance our knowledge of what factors are and are not involved in regulating the progression of epigenetic aging.

### Future Directions

The RheMacAge model has the potential to complement and expand aging research in primate populations. Our model can be used to test the effects of medical interventions intended to delay age-related physiological decline, such as caloric restriction, rapamycin administration, or other pharmacological treatments, where studies in macaques or baboons are ongoing (e.g., Tarandovskiy et al., 2020). It is also particularly well-suited to situations where environment and behavior intersect. For example, the model could be used to examine whether adversity experienced early in life is linked to late-life changes in the pace of biological aging. Such research could uncover molecular mechanisms that regulate how stressors become biologically embedded and may help identify health or behavioral variables that contribute to increased resiliency. Finally, our findings highlight the utility of the RheMacAge clock for disentangling when and to what extent social factors influence the pace of aging. The ability to apply the same predictive model across species facilitates comparative work, which in turn highlights how variation in social hierarchies translates into variation in their physiological correlates.

Excitingly, the RheMacAge clock comes online at a time when resources for studying epigenetic aging in nonhuman mammals are expanding more generally. For example, the HorvathMammalMethylChip has already been deployed to study the effects of hibernation on epigenetic aging in yellow-bellied marmots (Pinho et al., 2022), to examine postnatal development of the epigenome in opossums and other marsupials (Horvath et al., 2022), and to evaluate the potential lifespan-extending effects of partial cell reprogramming in a mouse model of premature aging (Browder et al., 2022). Our RheMacAge model, which takes a sliding-window approach, provides a useful alternative to the HorvathMammalMethylChip for researchers who wish to look at how DNA methylation varies across the genome more broadly, by coupling applications of the clock with differential or allele-specific methylation analyses. In mammals, RRBS datasets typically profile upwards of ~500,000 CpG sites after quality control, as opposed to the 38,000 CpG sites on the chip. Additionally, RRBS datasets are more likely to contain sites that are specific to the species of interest, as CpG sites that are not as tightly conserved are often the most interesting from an evolutionary perspective. Future work that calibrates the epigenetic clock using blood chemistry or other relevant biological measures of aging to predict physiological decline and/or mortality, as the human “next-generation” clocks have found success in doing (e.g., Belsky et al. 2020; Levine et al. 2018; Lu et al. 2019), has the potential to further extend the applicability of these models. Together, these methods contribute to an important and growing analytical toolkit for research in nonhuman primates.

## CONCLUSIONS

We have developed a method that overcomes a persistent barrier to comparative analyses using BS-seq datasets. Our approach is easy to implement and the generalizability of the resulting model enables cross-study comparison, as demonstrated by the successful application of our RheMacAge model to independent RRBS datasets to predict age in two distinct species. Our model recapitulated a previously identified relationship between rank and epigenetic aging in wild male baboons (Anderson et al., 2021) but found no such effect in male rhesus macaques, suggesting the importance of the way dominance hierarchies are formed and maintained in different species. These results provide proof-of-concept for our model and its capacity to measure the influence of the social and ecological environment on health, aging, disease and mortality risk. Despite numerous attempts to decipher the underlying mechanisms of the clock, they remain largely obscure. However, such knowledge is not required to use the clock to continue to probe environmental variables that accelerate or decelerate the pace of aging. The increasingly widespread use of epigenetic clock models has advanced the field towards an essential goal: quantification of the impact of lived experiences on health and aging. Future research should aim to identify specific variables that have the greatest impact on health and longevity. We anticipate that our model will facilitate interrogation of novel socio-environmental factors and whether they are or are *not* able to effectively “get under the skin” to engender epigenetic change, and how these modifications fit into the larger, systemic picture of aging and longevity across species.

## ACKNOWLEDGEMENTS

We thank the Caribbean Primate Research Center and staff from the Cayo Santiago Field Station, as well as Luis Barreiro, Mark Wilson, and the staff at the Yerkes National Primate Research Center. This research was supported by the National Science Foundation (BCS-1920350, BCS-1800558), and the National Institutes of Health (R21-AG075648, R00-AG051764, R01-AG060931, R01-MH096875, R01-MH089484, R01-MH118203, R01-GM102562, R01-AG057235, P51-OD011132). Cayo Santiago Field Station is supported by the Office of Research Infrastructure Programs [ORIP] of the NIH (P40-OD012217). K.L.C. (T32-AG000057) and M.M.W. (F31-AG072787) were supported during this work by National Institutes of Health fellowships.

This research was facilitated through the use of advanced computational, storage, and networking infrastructure provided by the mox computing cluster at the University of Washington, the Talapas computing cluster at the University of Oregon, and the Agave computing cluster at Arizona State University.

## DATA ACCESSIBILITY STATEMENT

Code for all analyses can be found on GitHub (https://github.com/elisabethgoldman/RheMacAge) and data are accessible on NCBI SRA through BioProject accession PRJNA610241.

## AUTHOR CONTRIBUTIONS

NSM, KNS, LJNB, JPH, and EAG conceptualized the research; Sample collection on Cayo Santiago was facilitated by KLC, JEH, MIM, MJM, JPH, LJNB, MLP, and NSM; KLC, AM, and SNS performed genomic lab work; behavioral data collection on Cayo Santiago was coordinated and led by LJNB; sample collection from the Yerkes Primate Research Center was facilitated by NSM and JT; JT oversaw data collection from the baboons at Amboseli; EAG, KLC, and MMW performed genomic analysis with input from NSM, JT, and JAA; EAG, KLC, MMW, KNS, and NSM wrote the manuscript. All authors reviewed and revised the manuscript.

