## Supplemental Figures for "A generalizable epigenetic clock captures aging in two nonhuman primates"

**CAYO BIOBANK RESEARCH UNIT**
Susan C. Antón, Lauren J. N. Brent, James P. Higham, Melween I. Martínez, Amanda D. Melin, Michael J. Montague, Michael L. Platt, Jérôme Sallet, and Noah Snyder-Mackler.

**SUPPLEMENTAL FIGURES**

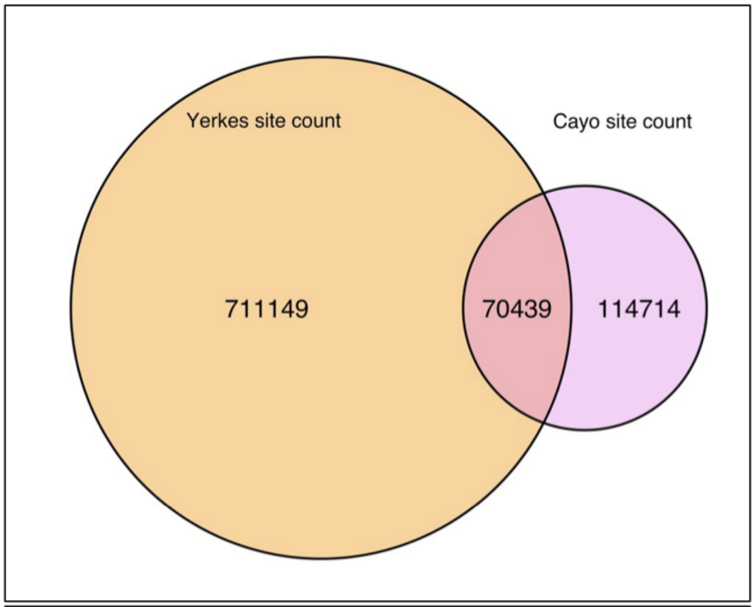


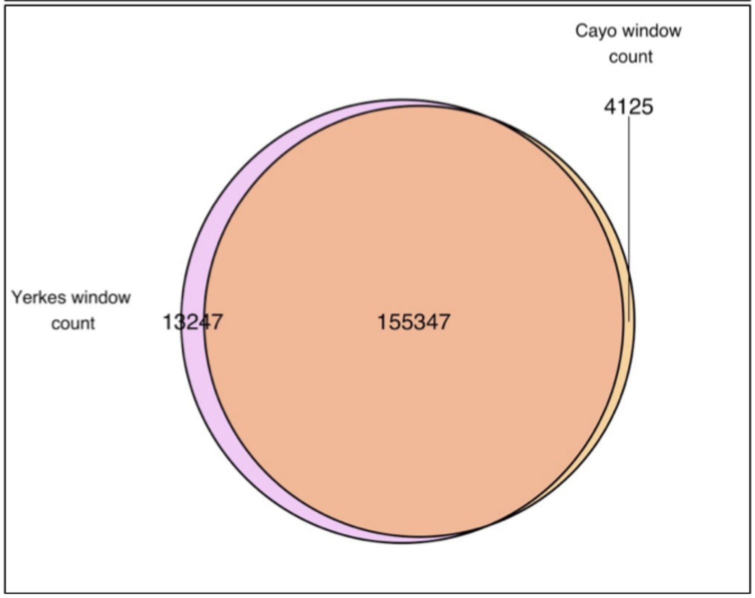


**Figure S1 Proportion of Shared Loci Between Site-Based and Window-Based Models**

The proportion of shared coverage between our rhesus training and test datasets increased from 38% for the site-based approach (top) to 97% for the window-based approach (bottom). Numbers on the left of each Venn diagram show the number of features (sites or windows) unique to the Yerkes dataset, while numbers on the right-hand side show those unique to the Cayo Santiago dataset. The numbers shown where the two circles overlap refer to the count of shared features between the two datasets when either approach was used.

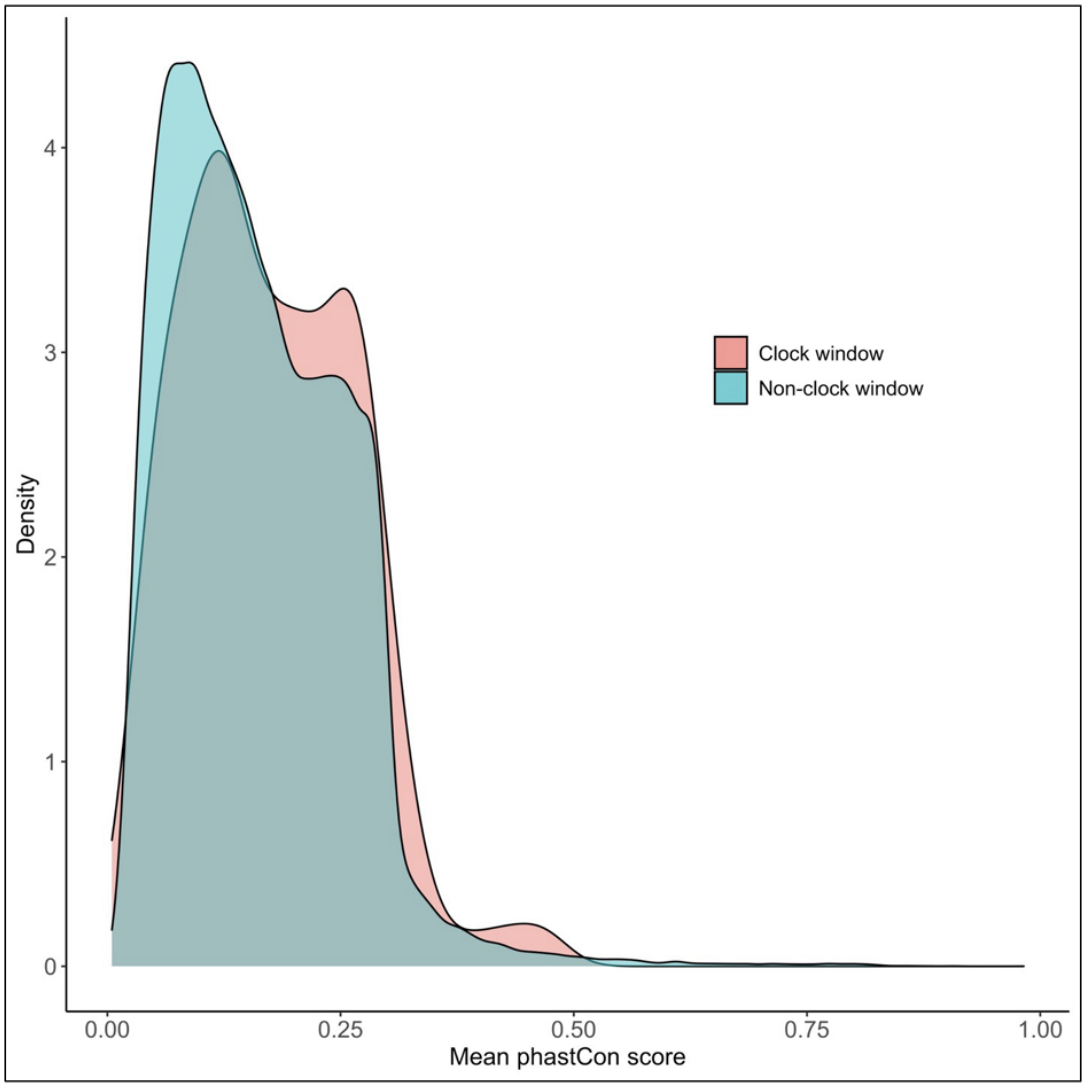


**Figure S2 Comparison of Enrichment for Conserved Sequences Between Clock and Non-Clock Windows**

Clock windows are modestly but significantly enriched for evolutionarily conserved sequences (two-sample Kolmogorov-Smirnov test, $D = 0.09$, $p = 0.007$).

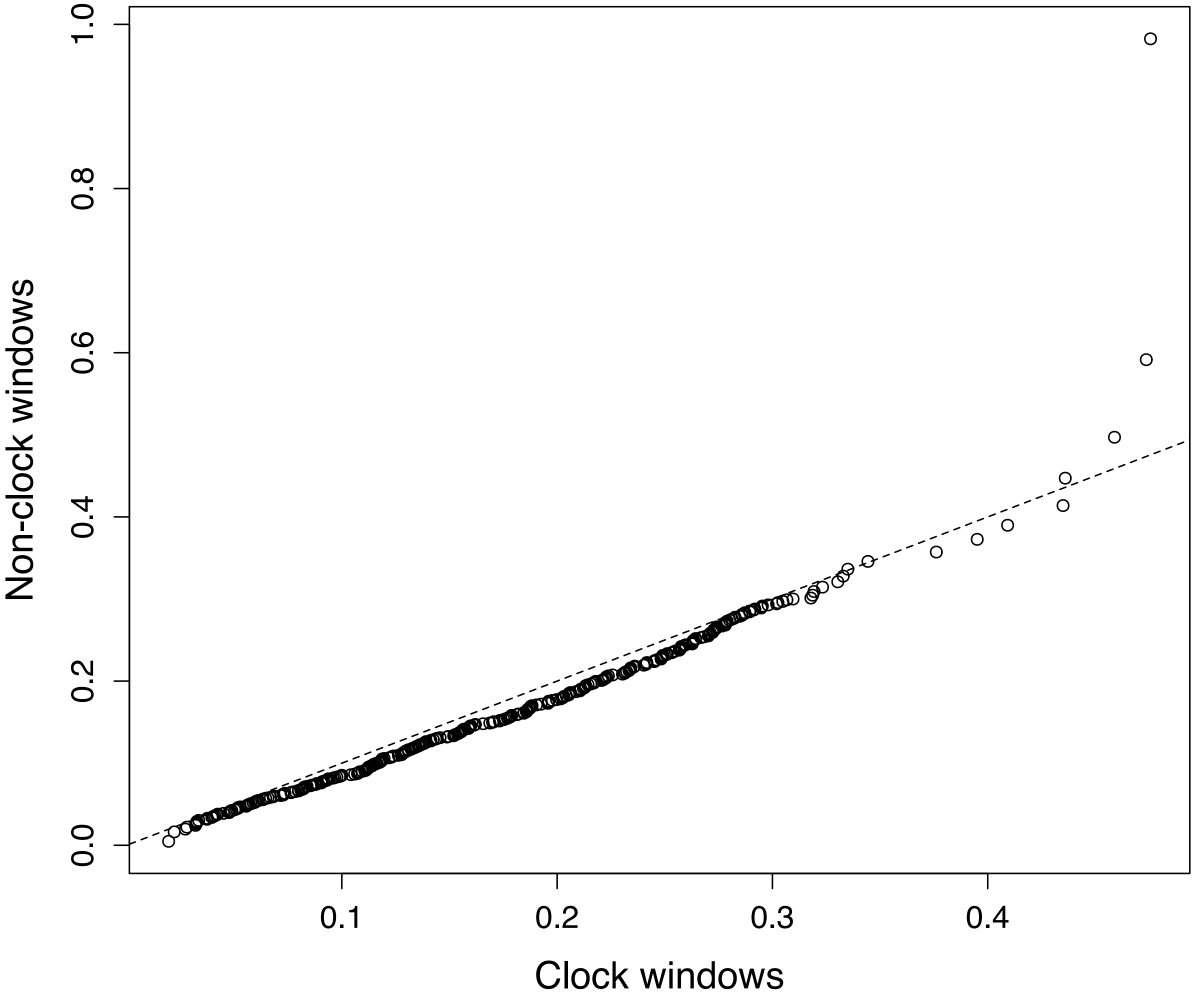


**Figure S3 Comparison of Sequence Conservation Between Clock and Non-Clock Windows**

Distribution of mean phastCon scores for clock windows (those automatically selected for inclusion in the model) and non-clock windows. Clock windows show a modest trend towards sequence conservation as compared to the windows that were not selected. Dashed line shows x=y.

### METHODS SUPPLEMENT

*Bismark Parameter Settings*

When aligning the reads to the reference genome using Bismark (Krueger & Andrews, 2011), we made two modifications to the alignment default parameters to reduce the number of ambiguously mapped reads (which result in data loss because these reads are discarded). We relaxed the minimum alignment score to allow approximately 3 mismatches or gaps in the alignment of 1-2 bp each (this is the "--score_min" parameter; note that the maximum alignment score is zero, corresponding to perfect alignment with no mismatches or gaps). We also increased the number of times Bismark attempted to reseed a repetitive (low complexity) read before marking it as invalid, from a default of 2 to 8 times ("-R" parameter).

*Criteria Used to Eliminate Samples from the Site-Based Dataset*

Our initial dataset contained 631 genomic libraries. We removed 21 low coverage libraries following alignment to the reference genome. We combined data from duplicate libraries (those derived from samples collected from the same individual on the same day, n = 29), leaving 581 samples. We removed eight samples after plotting the ratio of X-chromosome to chromosome 19-mapping sites by sex and finding four samples labeled as female that clustered with the males, and four labeled as male that clustered with the females, suggesting they had been mislabeled. Finally, we removed 24 samples that were missing > 25% of their data in the final filtered dataset.

*Considerations for Training, Optimizing, and Implementing Epigenetic Clock Models*

We used an elastic net penalized regression algorithm that automatically selects different subsets of CpG sites (or 1 kb windows) that together generate the most accurate age predictions. We used a nested loop structure to train and optimize our penalized regression model, with a leave-one-out cross validation (LOOCV) outer loop to tune model hyperparameters and an inner 10-fold cross validation loop to fit the model to training data, which determines the model's coefficients.

In the case of DNA methylation-based epigenetic clocks, our goal is to model the relationship between methylation at CpG sites or windows (independent variables) and chronological age (dependent variable).

We first fit a model to our training dataset. During the training stage, the algorithm is given both the methylation ratios and the chronological age of each sample. The algorithm then uses N-1 samples to train "proto-models" by dividing the data into 10 folds and running an internal cross-validation loop to train and validate on the inner folds. It determines which combination of features predict calendar age while minimizing the mean squared error. Hyperparameters are meta-parameters that are not learnable from the training data; examples are regularization parameters like lambda or the value of *K* in K-fold cross validation. We initially use previous knowledge or default settings for the hyperparameters and can subsequently optimize them by using caret's train() function and setting different hyperparameter combinations using preProcOptions(). Tuning alpha may only result in modest boosts in performance, but it is still recommended to examine results from setting different alpha values during the model optimization process.

*Enrichment Analysis for Evolutionarily Conserved Sequences*

To assign conservation scores to windows in rhesus macaque coordinates, we calculated phastCons scores directly using the "57 mammals EPO" multiple species alignment obtained from Ensembl (release 101). Multiple alignment format (MAF) files were processed in the following manner: First, ancestral species were removed, along with blocks not containing rhesus macaque sequences using maffilter v1.3.1 (Dutheil et al., 2014). We then removed species duplicates from each alignment and indexed each block to the rhesus macaque reference genome using mafTools (Mayakonda et al., 2018). Next, we used maf_parse from the PHAST utilities (Hubisz et al., 2011) to extract blocks corresponding to the 155,347 windows in this analysis. After extracting each window, we performed a local realignment of each block using MAFFT (v7.402) (Katoh & Standley, 2013) and maffilter. Some rhesus sequences that were originally from the same window were split across multiple blocks or MAF files. We thus rearranged alignment blocks such that each MAF file contained only blocks from the same rhesus macaque window using a custom shell script. We then combined blocks using the Merge() function in maffilter with rhesus macaque set as the reference species. We calculated conservation scores using the phastCons program (v1.5). First, we fit a phylogenetic model using phyloFit, the REV nucleotide substitution model, and the phylogenetic tree provided with the dataset in the Ensembl release. A minority of windows were excluded (7,008, or 4.5%) from this analysis because they were not represented in the multiple species alignment. We ran phastCons using the arguments "--expected-length 45 --target-coverage 0.3 --rho 0.3", which are identical to arguments used in the UCSC Genome Browser pipeline for generating conservation tracks (e.g., http://genome.ucsc.edu/cgi-bin/hgTrackUi?g=cons100way). The resulting phastCons scores represent probabilities of negative selection at the per-site level. We summarized each window by calculating the mean phastCons score across all windows.

*Sample Removal in the Test Datasets*

For the Yerkes macaques, we removed two samples due to insufficient library size (remaining n = 43). For the Amboseli baboons, we removed nine samples that failed three attempts at alignment to the bisulfite-converted rhesus genome (Mmul10), and six with low library sizes (remaining n = 271).
